# Advancements in Inflammation Parallels Myopenia in *Winnie* Mice Model of Spontaneous Chronic Colitis

**DOI:** 10.1101/2024.11.20.624215

**Authors:** Shilpa Sharma, Danielle Debruin, Jeannie Devereaux, Kulmira Nurgali, Alan Hayes, Gustavo Duque

## Abstract

**Background:** Inflammatory bowel disease (IBD) is characterized by gastrointestinal inflammation and systemic complications, including muscle wasting. Musculoskeletal conditions have remained an unappreciated aspect of IBD. This study aimed to describe the Winnie mouse model’s skeletal muscle phenotype and functional alterations. This model spontaneously develops chronic colitis that closely resembles human IBD.

**Methods:** *Winnie* mice and C57BL/6 (littermate) controls were evaluated at 5-6 weeks (pre-colitis) and 15-16 weeks (active colitis). Assessments included disease activity, inflammation, muscle function tests, and *ex vivo* analyses of the soleus (SOL) and tibialis anterior (TA) muscles for mass and histology.

**Results:** At 5-6 weeks, *Winnie* mice showed no disease activity or muscle changes. At 15 weeks, *Winnie* mice exhibited significantly higher disease activity index (DAI) and elevated lipocalin-2 (LCN-2) levels than controls. SOL and TA muscles showed decreased weights and fibre sizes, which correlated significantly with LCN-2 levels. Wheel-running activity was reduced, which correlated with increased LCN-2 levels and DAI. DAI negatively correlated with muscle mass and fibre size. Grip strength remained unaltered despite these changes.

**Conclusion:** The *Winnie* mouse model develops significant skeletal muscle alterations paralleling intestinal inflammation progression, characterized by reduced muscle mass and size with preserved strength but impaired functional endurance capacity. These findings establish the *Winnie* model as a valuable tool for investigating IBD-associated muscle wasting.

**Key Messages:** **1. What is already known?**

- Inflammatory bowel disease (IBD) is associated with systemic complications, including muscle dysfunction.

**2. What is new here?**

- The *Winnie* mouse model of chronic colitis develops skeletal muscle alterations that correlate with the progression of intestinal inflammation.
- Muscle changes in *Winnie* mice include reduced muscle mass, decreased fibre size, and, functional decline but preserved grip strength.

**3. How can this study help patient care?**

- This research highlights the importance of monitoring muscle health in IBD patients and suggests potential targets for interventions to preserve muscle function.
- The *Winnie* model provides a valuable tool for testing therapies aimed at addressing muscle dysfunction in IBD.

## Introduction

Inflammatory bowel disease (IBD) is recognized as a complex, chronic disorder primarily affecting the gastrointestinal tract (GIT). IBD primarily comprises two chronic idiopathic conditions: ulcerative colitis (UC) and Crohn’s disease (CD). These disorders exhibit distinct inflammatory patterns within the GIT. UC is characterized by mucosal and submucosal ulcerations confined to the rectum and colon. In contrast, the CD can affect any part of the GIT, featuring transmural inflammation, skip lesions, and potential stricture formation.^1^

While IBD primarily manifests as gut pathology, it often entails systemic complications.^2^ One such complication is the progressive, generalized loss of muscle mass and function, contributing to increased morbidity in IBD patients.^5,6^ A recent study by Fatani et al. (2023) revealed alarming statistics: 42% of adult IBD patients exhibit myopenia (low muscle mass), 34% demonstrate pre-sarcopenia (low muscle strength), and 17% present with sarcopenia (low muscle mass and strength). The prevalence is particularly high in females (54%) compared to males (45%).^3^ Sarcopenia can lead to an increased risk of falls, fractures, and fatigue perception, affecting patients’ mobility, independence, and overall quality of life.^8^ Despite the clinical significance of muscle wasting in IBD, the underlying mechanisms remain poorly understood. The etiology of sarcopenia concomitant with IBD is multifactorial, involving persistent inflammation, physical inactivity, malnutrition, and iatrogenic factors such as corticosteroids use. However, the exact pathophysiology of sarcopenia in IBD remains unknown.^4,5^

To address these knowledge gaps, animal models provide a valuable platform for investigating the underlying molecular and cellular processes that contribute to muscle atrophy and dysfunction in chronic inflammation. While chemically-induced colitis models using dextran sodium sulfate (DSS) and trinitrobenzene sulfonic acid (TNBS) are widely used due to their accessibility, low cost, and rapid development, they fail to fully capture the complexity of human IBD.^6^ In contrast, spontaneous colitis models offer invaluable insights for developing novel therapeutic interventions, as they more closely mimic IBD pathogenesis. Two notable spontaneous colitis models are: SAMP1/SkuSlc mouse model, characterized by ileal lesions, which closely resembles active CD^7^ and *Winnie* mouse model of spontaneous chronic colitis, which primarily affects the large intestine, closely mirrors UC.^8,9^

The *Winnie* mouse model develops spontaneous colitis due to a missense mutation in the *Muc2* mucin gene, without the need for experimental intervention. This genetic basis for colitis development contrasts with other experimental models that require chemical induction or immune cell transfer.^10–12^ The mutation leads to endoplasmic reticulum stress in goblet cells and subsequent mucosal damage.^9,13^ Defects in the protective mucin layer increase intestinal susceptibility to antigens, resulting in an abrupt immunological response, microbiota dysbiosis, and alterations in GIT function, characterized by an inflammatory cytokine profile similar to UC.^14–16^ The chronic nature of the disease in *Winnie* mice provides a unique opportunity to investigate long-term morphological and physiological changes in relevant tissues, including potential musculoskeletal complications.

This study aims to characterize the structural and functional changes in skeletal muscle associated with chronic colitis using the *Winnie* mouse model. By examining these changes, we seek to elucidate the relationship between chronic intestinal inflammation and muscle health, potentially informing future therapeutic strategies for IBD-associated muscle wasting. We hypothesized that *Winnie* mice, a model of chronic colitis, would exhibit alterations in muscle phenotype and function similar to those observed in UC-associated sarcopenia.

Sarcopenia, characterized by progressive loss of skeletal muscle mass, strength, and function, has been documented in patients with chronic UC.^17^ By investigating these muscle changes in the *Winnie* mouse model, we aimed to elucidate the relationship between chronic intestinal inflammation and skeletal muscle alterations, potentially mirroring the muscle wasting observed in UC patients. To test this hypothesis, we conducted a comprehensive analysis of muscle phenotype and function in *Winnie* mice at 5–6 weeks (pre-inflammatory stage) and 15–16 weeks (active colitis stage), comparing them to age- and sex-matched C57BL/6 littermate controls. Our investigation focused on two key muscle groups: the soleus (SOL), a predominantly slow-twitch muscle, and the tibialis anterior (TA), a predominantly fast-twitch muscle. We assessed multiple parameters including muscle weights, muscle fibre size, physical activity, and grip strength. By examining both *ex vivo* muscle properties and *in vivo* functional tests like physical activity and grip strength, our goal was to address gaps and provide a holistic view of muscle health in the context of chronic colitis.

## Materials and Methods

### Animals

*Winnie* mice (Muc2 mutant) were obtained from the Victoria University animal facility (Melbourne, VIC, Australia), while wild-type C57BL/6 mice were sourced from the Animal Resources Centre (Perth, Western Australia). All mice were given at least 1 week of acclimatization at the Australian Institute for Musculoskeletal Science (AIMSS) animal facility (Melbourne, Victoria, Australia). These mice strains were used to generate heterozygous *Winnie*^+/-^ breeders. To establish our experimental cohorts, these breeders were co-housed to obtain *Winnie*^-/-^ mice and their control C57BL/6 littermates from the same mother. All mice were co-housed to ensure exposure to comparable mucosal environments. For our experiments, we utilized a total of 60 mice: 33 *Winnie*^-/-^ and age- and sex-matched 30 control C57BL/6 controls. Mice were studied at two age groups: 5–6 weeks and 15–16 weeks. Animals were housed in a climate-controlled room maintained at 23 ± 2°C and 55 ± 5% relative humidity, with a 12-hour light/dark cycle. Standard laboratory chow and water were provided *ad libitum*. We ensured consistent environmental conditions (e.g., cage location, diet, water, and housing conditions) for both *Winnie* and control groups within our animal breeding and holding facility. Animal welfare was closely monitored throughout the study. In cases where mice exhibited severe illness, including signs of significant bleeding, were promptly and humanely euthanized immediately to prevent unnecessary suffering in line with established ethical protocols for animal research. All procedures were conducted in accordance with the Australian code of practice for the care and use of animals for scientific purposes and were approved by the Victoria University Animal Experimentation Ethics Committee (AEEC17/016).

### Colitis severity assessment

Colitis severity in *Winnie* mice was characterized by parameters, such as changes in body weight, stool consistency, and rectal examination compared to C57BL/6 controls at 5 and 15 weeks of age. Changes in body weight were calculated as a percentage difference from the baseline weights of 5-week-old C57BL/6 controls. Stool consistency (normal, loose, or diarrheal) was assessed visually and assigned a score. The rectal examination involved inspection of the perianal region and a score was assigned based on the presence and severity of rectal bleeding. Scoring of disease parameters in *Winnie* mice compared to age-matched C57BL/6 controls was done based on the percentage of weight loss (e.g., 0 = no weight loss, 1 = 1-5% weight loss, 2 = 5-10% weight loss, 3 = 10-20% weight loss, 4 = >20% weight loss), stool consistency (0 = normal, 1 = loose stools, 2 = diarrhea), and rectal bleeding (0 = no blood, 1 = bleeding, 2 = rectal prolapse). The DAI is calculated as the sum of the scores for weight loss, stool consistency, and rectal bleeding.

### Faecal Lipocalin-2 assessment

To assess intestinal inflammation in *Winnie* mice, fecal LCN-2, a highly sensitive biomarker for intestinal inflammation, was measured.^18^ Fecal pellets were homogenized in phosphate-buffered saline (PBS) containing 0.1% Tween 20 at a concentration of 100 mg/mL. The homogenates were centrifuged for 10 minutes at 12,000 rpm and 4°C. LCN-2 was quantified in the clear supernatants using a mouse NGAL ELISA kit (LCN-2, ab119601).^19^ A microplate reader measured absorbance at 450 nm, with a correction wavelength of 540 nm, to determine LCN-2 protein concentration (ng/mL) in the fecal supernatants.

### Muscle collection and histological analysis

SOL and TA muscles were immediately excised, weighed, and embedded in an optimal cutting temperature (OCT) compound. They were then snap-frozen in partially thawed isopentane (Sigma Aldrich, Castle Hill NSW, Australia) and stored at −80°C until histological analysis. Transverse muscle sections (12 µm thick) were cut from the mid-belly region of each muscle in a cryotome at −15°C and then air-dried for 30 min. For hematoxylin and eosin (H&E) staining, cryosections were dehydrated by sequential immersion in ethanol gradients (100%, 95%, 70%) for 2 minutes each. Samples were briefly rinsed in tap water for 30 seconds. For nuclear staining, samples were immersed in hematoxylin (Sigma-Aldrich) for 1 minute, followed by a tap water rinse. After immersion in Scott’s tap water for 1 minute, cytoplasmic staining was performed by immersing the sections in eosin (Sigma-Aldrich) for 3 minutes, followed by a rinse with tap water. Then, the sections were dehydrated by immersion in 100% ethanol for 1 minute, followed by clearing in xylene for 5 minutes. Finally, sections were mounted on glass slides using a distyrene-plasticiser-xylene (DPX) mounting medium (BDH, Poole, UK). Muscle cross-sections were visualized using a Zeiss Axio Imager Z2 microscope (Carl Zeiss Microimaging GmbH, Oberkochen, Germany) at 20× magnification. Images were acquired using the MetaSystems Metaphor program. Subsequently, these images were digitally stitched together to create composite views of the entire muscle cross-section using VSlide software. Muscle fibre size was quantified using the minimal Feret’s diameter method, analyzing 200-250 fibres per muscle. All histological analyses were conducted in a blinded fashion to minimize bias. Minimal Feret’s diameters were measured and recorded using ImageJ software (National Institutes of Health, Bethesda, MD, USA), with measurements calibrated against a stage micrometer to ensure accuracy. Minimal Feret’s diameter represents the minimum distance between two parallel tangents at opposing borders of the muscle fibre, regardless of orientation. This method was chosen for its robustness against variations in tissue sectioning angle, which can significantly affect other size parameters.^20^

### Physical activity analysis using metabolic cages

An eight-channel Promethion system (Sable Systems International) was used. Mice were individually housed with ad libitum access to food, water, and a running wheel in Promethion metabolic cages. X-Y- and Z-axes photoelectric beam motion detectors were also positioned around the cage. An initial 4-day acclimatization period allowed mice to adjust to the new environment and establish baseline running activity. Data presented are from the final 24 hours of recording, divided into diurnal (7 am to 7 pm) and nocturnal (7 pm to 7 am) phases. Data points were averaged into 4 minute intervals using ExpeData PRO Software with a customized macro 6 provided by Sable Systems.

### Grip strength analysis

Muscle strength of the forelimbs of *Winnie* mice at 15 weeks compared to age- and sex-matched C57BL/6 mice were measured using a grip strength test as previously described.^21^ Briefly, a grip test apparatus was used, which consisted of a stainless-steel wire mesh. Mice were held by the base of their tail and allowed to grasp the grip test apparatus with their forepaws. After the mouse held the mesh, the mouse’s tail was pulled at a constant slow speed to permit the mouse to build up resistance against it, until its grasp was broken. The transducer saves the value at this point. The test was repeated 3-5 times to obtain the best performance. A 30 s rest was used between lifts. Measurements of force exerted were collected for the mice and were compiled and analysed.

### Statistical Analysis

Statistical analyses were performed using GraphPad Prism v9.3 (GraphPad Software Inc., San Diego, CA, USA) and Microsoft Excel 2019 (Microsoft Corporation, Redmond, WA, USA). Before applying the statistical significance tests, a Shapiro-Wilk test was used to evaluate the normality of the data distribution. When the data sets had a normal distribution, t-tests were employed. A *P* value of < 0.05 was considered statistically significant. For unnormalized data, the non-parametric Mann–Whitney U-test was used to compare differences between the two groups. All data were expressed as mean ± SEM. Relationships between X and Y variables were assessed using linear regression analysis. *P* values for significant relationships were recorded. Additionally, the coefficient of determination (R²) was calculated and recorded to quantify the proportion of variance in the dependent variable that is predictable from the independent variable.

## Results

### Winnie mice with chronic colitis exhibit increased disease activity and elevated intestinal inflammation

At 5 weeks of age, no significant differences were observed between *Winnie* mice and C57BL/6 controls in any of the measured parameters (body weight, stool consistency, rectal bleeding). However, by 15 weeks of age, *Winnie* mice exhibited these symptoms, indicative of colitis. *Winnie* mice produced loose stools compared to well-formed faecal pellets in control mice at 15 weeks. The image showing prominent rectal prolapse in the *Winnie* mouse, indicated by the protruding rectal tissue is shown in **Supplementary Fig. 1**. This feature is absent in the C57BL/6 control mouse, which displays a normal anal region. No significant differences were observed in DAI scores between *Winnie* mice and controls at 5 weeks of age. DAI scores were significantly higher in *Winnie* mice compared to C57BL/6 controls at 15 weeks of age in both females (*P* < 0.0001) and males (*P* < 0.0001) (**Fig. 1A, 1A’**). Faecal LCN-2 levels, a robust marker of intestinal inflammation, were similar in 5-week-old *Winnie* mice and age- and sex-matched C57BL/6 controls. However, LCN-2 levels were elevated in 15-week-old *Winnie* mice compared to age- and sex-matched C57BL/6 controls (females: *P* < 0.0001; males: *P* < 0.001) (**Fig. 1B, 1B’**).

**Figure 1.**
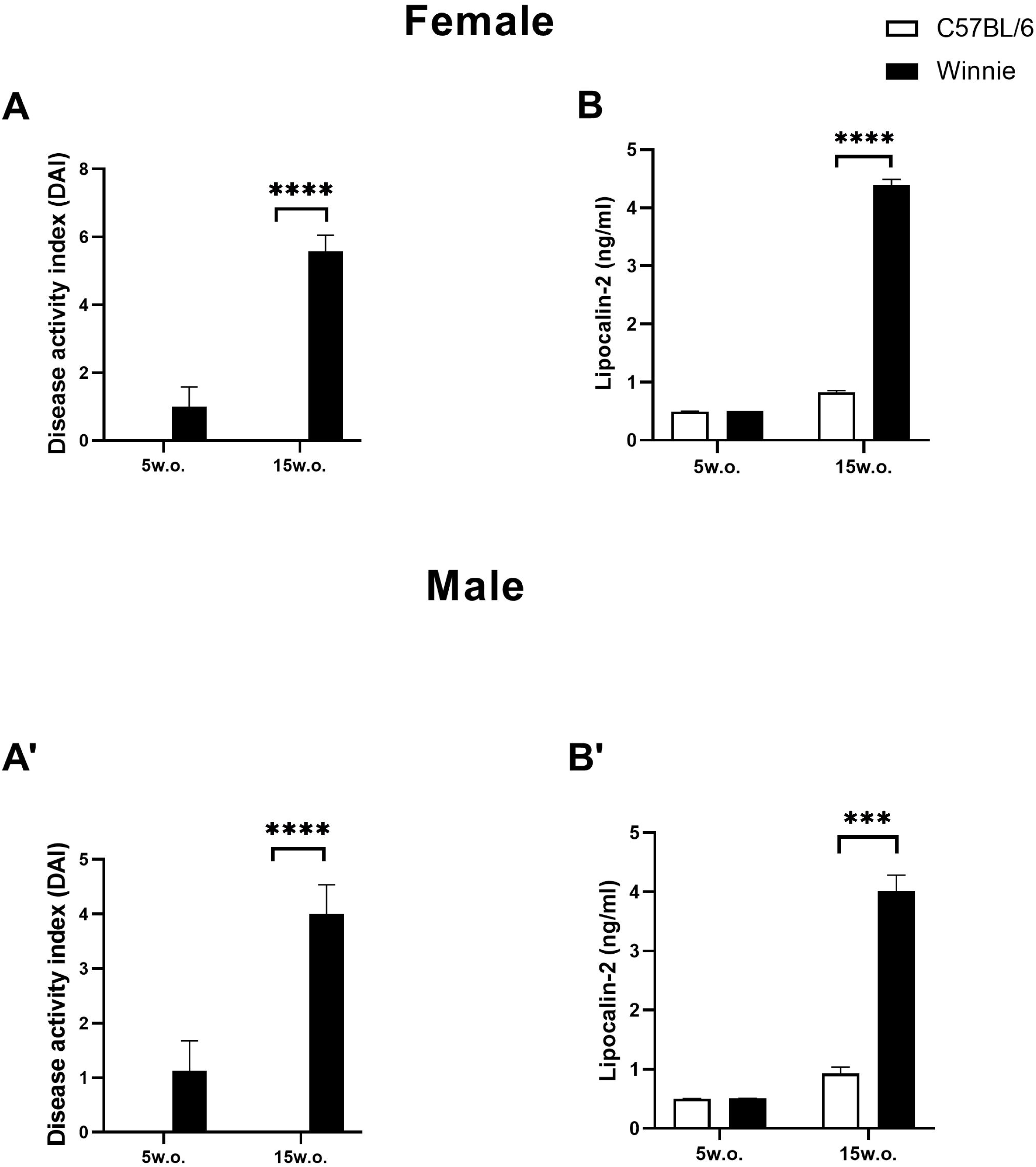
Progression of colitis symptoms and associated changes of C57BL/6 and *Winnie* mice at 5 and 15 weeks of age. (**A, A’**) Disease activity index from female and male *Winnie* mice at 5 and 15 weeks compared to C57BL/6 controls. (**B**) Faecal lipocalin-2 levels of *Winnie* mice compared to age- and sex-matched C57BL/6 controls. (**B, B’**) Lipocalin-2 levels were quantified from faecal samples of female and male *Winnie* mice compared to age- and sex-matched C57BL/6 controls at 5 and 15 weeks of age. w.o. = weeks old. Data are presented as mean ± SEM. *P < 0.05, **P < 0.01; ****P < 0.0001; n=6-8 mice/group mice/group.

### Decrease in body weight, lean mass, and muscle mass of slow-twitch (SOL) and fast-twitch (TA) muscles in Winnie mice at 15 weeks of age

At 5 weeks of age, both female and male *Winnie* mice exhibited body weights comparable to age- and sex-matched C57BL/6 controls. However, by 15 weeks of age, *Winnie* mice of both sexes demonstrated significantly lower body weights compared to controls (*P* < 0.01 for both females and males) (**Fig. 2A, 2A’**). Preliminary observations using EchoMRI analysis revealed reduced lean mass (*P* < 0.01) and no change in fat mass in 15-week-old *Winnie* mice compared to age-matched C57BL/6 controls (**Supplementary Fig. 2A, 2B**). At 5 weeks of age, no significant differences were observed in muscle mass between *Winnie* mice and C57BL/6 controls for both SOL and TA muscles. However, by 15 weeks of age, absolute SOL muscle mass was decreased (*P* < 0.001 for both sexes) (**Fig. 2B, 2B’**). Analysis of SOL muscle mass normalized to body weight showed a reduction (females: *P* < 0.05, males: *P* < 0.001) (**Fig. 2C, 2C’**). Absolute TA muscle mass was decreased (females: *P* < 0.05; males: *P* < 0.001) (**Fig. 2D, 2D’**). Analysis of TA muscle mass normalized to body weight showed a reduction (*P* < 0.01 for both sexes) (**Fig. 2E, 2E’**).

**Figure 2.**
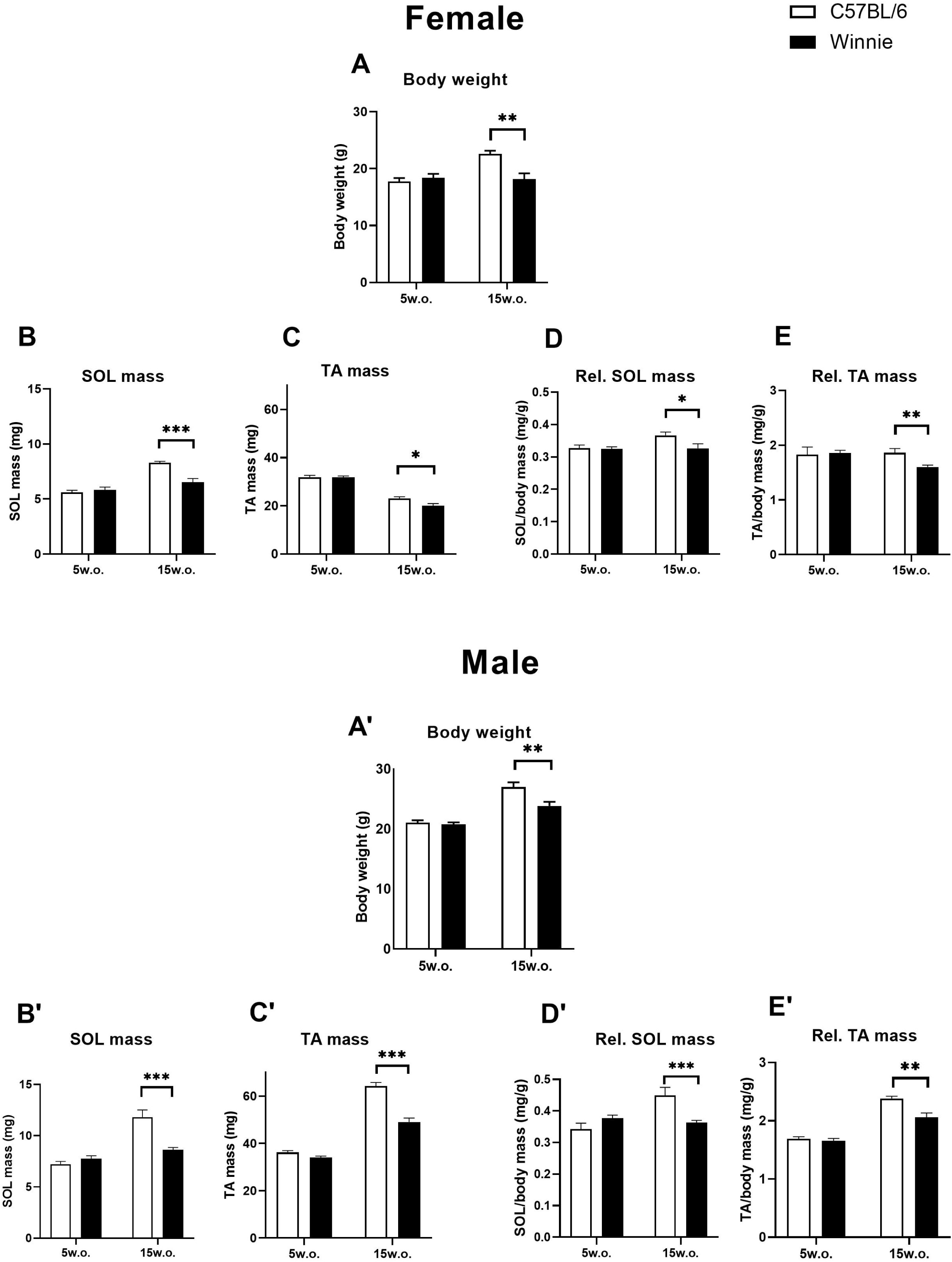
Changes in body weight and muscle mass of soleus (SOL) and tibialis anterior (TA) muscles in C57BL/6 and *Winnie* mice at 5 and 15 weeks of age. (**A, A’**) Body weight of female and male Winnie mice compared to C57BL/6 control mice at 5 and 15 weeks. (**B, B’**) Absolute SOL mass in females and males, respectively. (**C, C’**) SOL mass normalized to body mass in females and males, respectively. (**D, D’**) Absolute TA mass in females and males, respectively. (**E, E’**) Absolute and relative mass of SOL muscle and TA mass normalized to body mass in females and males, respectively. w.o. = weeks old. Data is represented as a mean ± SEM. *P < 0.05, **P < 0.01, ***P < 0.001, n=5-8 mice/group.

### Muscle-specific alterations in fibre size distribution in the Winnie mouse model of colitis

At 5 weeks of age, no significant differences in muscle fibre size were observed between *Winnie* and C57BL/6 mice for both SOL and TA muscles in either sex. By 15 weeks of age, minimal Feret’s diameter of SOL and TA were significantly decreased (females: *P* < 0.01, males: *P* < 0.01 for SOL; females: *P* < 0.001, males: *P* < 0.05 for TA) (**Fig. 3A-3D, 3A’-3D’**). The fibre size distribution in both muscles shifted towards smaller diameters, indicating a reduction in the proportion of larger fibres and an increase in smaller fibres. Importantly, these alterations in muscle fibre size were not observed in 5-week-old *Winnie* mice, which displayed fibre size distributions similar to age- and sex-matched C57BL/6 controls in both SOL and TA muscles (**Fig. 3E-3H, 3E’-3H’**).

**Figure 3.**
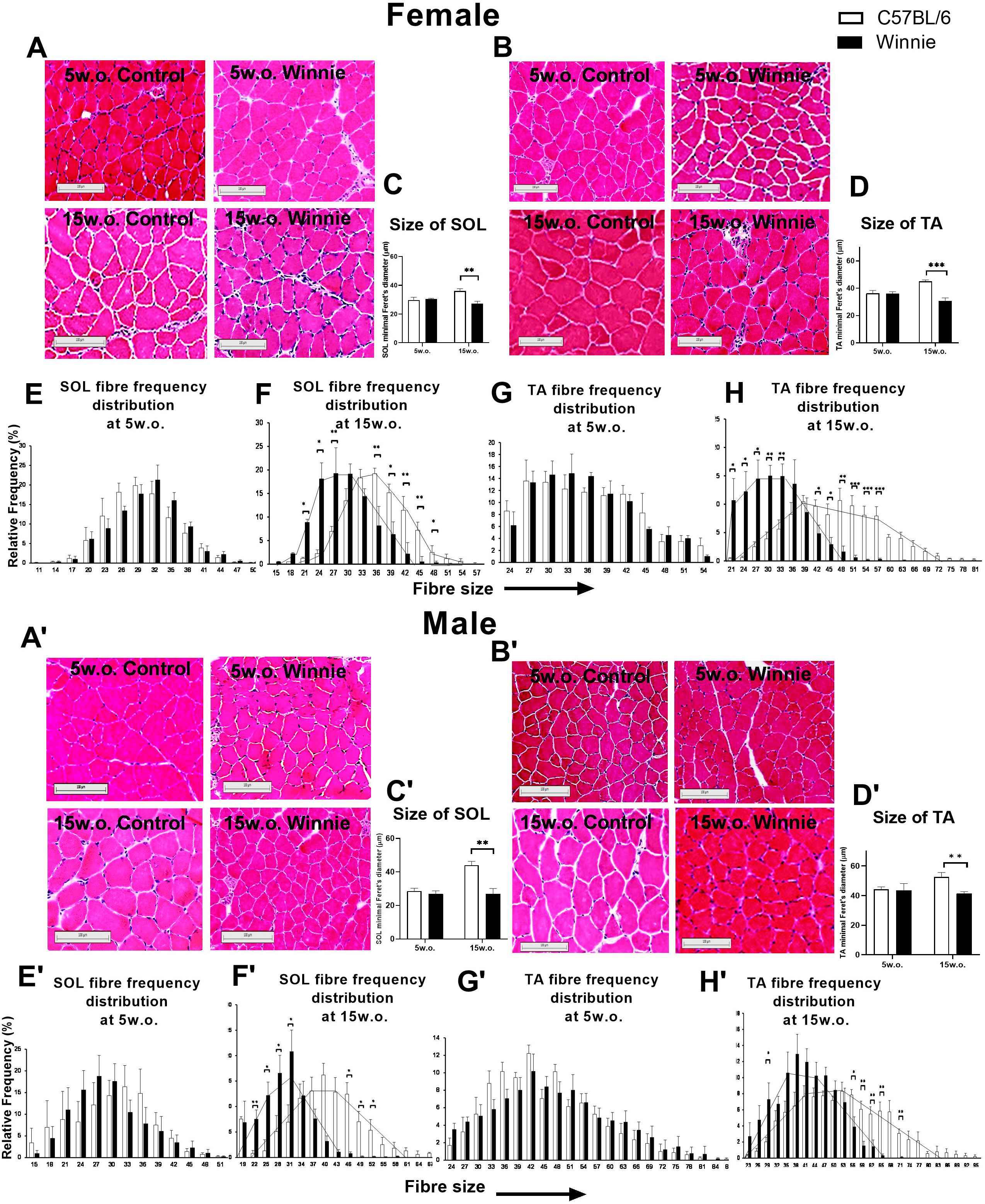
Analysis of muscle fibre size of soleus (SOL) and tibialis anterior (TA) muscles of C57BL/6 and *Winnie* mice at 5 and 15 weeks of age. (**A, A’, B, B’**) Representative images of SOL and TA muscle cross-sections at 5 and 15 weeks old for both genotypes and sexes. (**C, C’**) Minimal Feret’s diameter of SOL fibres of female and male *Winnie* mice compared to C57BL/6 controls at 5 and 15 weeks of age. (**D, D’**) Minimal Feret’s diameter of TA fibres of female and male *Winnie* mice compared to C57BL/6 controls at 5 and 15 weeks of age. Scale bars represent 100 μm. (**E, E’**) SOL fibre size frequency distribution female and male *Winnie* mice compared to C57BL/6 controls at 5 weeks. (**F, F’**) SOL fibre size frequency distribution old of female and male *Winnie* mice compared to C57BL/6 controls at 15 weeks. (**G, G’**) TA fibre size frequency distribution female and male *Winnie* mice compared to C57BL/6 controls at 5 weeks. (**H, H’**) TA fibre size frequency distribution of female and male *Winnie* mice compared to C57BL/6 controls at 15 weeks. Data are presented as mean ± SEM for Feret’s diameter and as relative frequency (%) for distribution graphs. *P <0.05, **P < 0.01, ***P < 0.001; n=5 mice/group.

### Association between intestinal inflammation, DAI, and skeletal muscle atrophy in Winnie mice with chronic colitis

Correlation analysis revealed negative associations between fecal LCN-2 levels and muscle mass as well as fibre size in both SOL and TA muscles. In the SOL muscle, negative correlations were observed with muscle mass (females: R^2^=0.572, *P* < 0.01; males: R^2^=0.5634, *P* < 0.01) and fibre size (females: R^2^=0.6844, *P* < 0.01; males: R^2^=0.6336, *P* < 0.01) (**Fig. 4A-4D**). Similarly, for the TA muscle, negative correlations were found with muscle mass (females: R^2^=0.7201, *P* < 0.0001; males: R^2^=0.6786, *P* < 0.001) and fibre size (females: R^2^=0.8303, *P* < 0.001; males: R^2^=0.747, *P* < 0.01) (**Fig. 4A’-4D’**).

**Figure 4.**
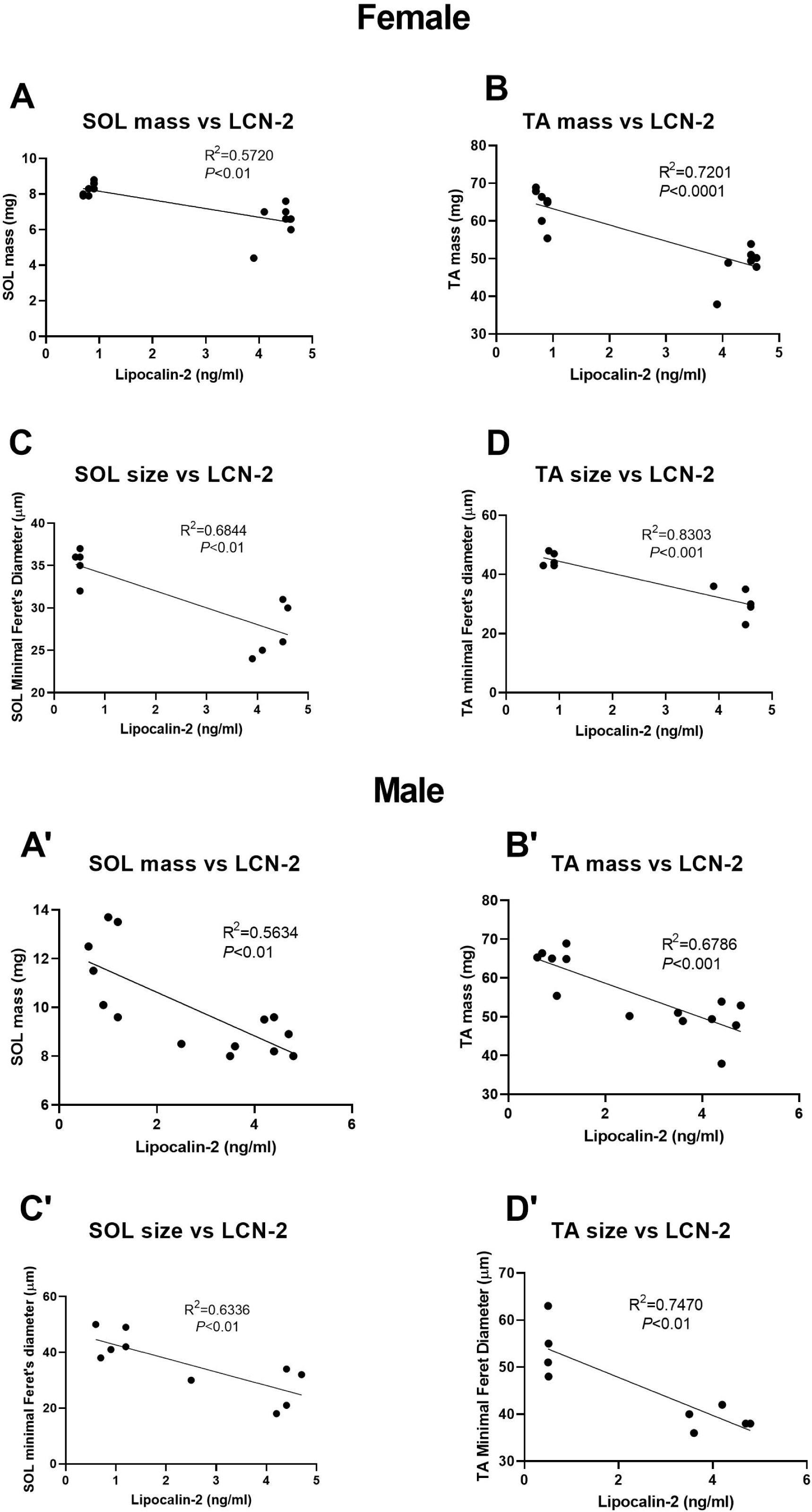
Linear regression analysis of faecal Lipocalin-2 (LCN-2) levels and muscle parameters (mass and size) of *Winnie* mice compared to C57BL/6 controls at 15 weeks. (**A, A’**) Correlations between LCN-2 levels and SOL muscle mass in females and males, respectively. (**B, B’**) Correlations between LCN-2 levels and TA muscle mass in females and males, respectively. (**C, C’**) Correlations between LCN-2 levels and SOL muscle fibre size in females and males, respectively. (**D, D’**) Correlations between LCN-2 levels and TA muscle fibre size in females and males, respectively. **P < 0.01, ***P < 0.001; n=9-14 mice/group. R^2^: coefficient of determination.

All correlations between DAI and muscle mass, as well as fibre size in both SOL and TA muscles were negative, indicating that higher disease activity is associated with lower muscle mass and smaller fibre diameters. For SOL, negative correlations were observed with muscle mass (females: R^2^ = 0.6505, *P* < 0.001; males: R^2^ = 0.5108, *P* < 0.01) and fibre size (females: R^2^ = 0.6115, *P* < 0.01; males: R^2^ = 0.7300, *P* < 0.01) (**Supplementary Figure 3A-3D**). In the TA muscle, negative correlations were observed with muscle mass (females: R^2^ = 0.3514, *P* < 0.05; males: R^2^ = 0.4315, *P* < 0.01) and fibre size (females: R^2^ = 0.5277, *P* < 0.05; males R^2^ = 0.4759, *P* < 0.05) (**Supplementary Figure 3A’-3D’**).

### Reduced wheel running activity but preserved grip strength in the Winnie mouse model of chronic colitis

Analysis of physical performance in 15-week-old *Winnie* mice revealed impairments compared to age- and sex-matched C57BL/6 controls, particularly during the nocturnal phase when mice are typically most active. Wheel running activity was markedly reduced in *Winnie* mice. Specifically, average running distance, speed, and time spent on the wheel were all decreased in 15-week-old *Winnie* mice compared to age- and sex-matched C57BL/6 controls (females: *P* < 0.001, *P* < 0.01, *P* < 0.05, respectively; males: *P* < 0.001, *P* < 0.0001, *P* < 0.01 respectively). During the diurnal phase, 15-week-old male *Winnie* mice exhibited decreased running distance, speed, and time spent compared to age- and sex-matched C57BL/6 controls (*P* < 0.01, *P* < 0.001, and *P* < 0.05, respectively), whereas in 15-week-old female *Winnie* mice, only running speed was decreased compared to age- and sex-matched C57BL/6 controls (*P* < 0.01) (**Fig. 5A-5C, 5A’-5C’**). Male *Winnie* mice showed decreased X-axis horizontal activity at 15 weeks compared to age- and sex-matched C57BL/6 controls at both diurnal and nocturnal phases (*P* < 0.05) (**Fig. 5D, 5D’**). No changes were observed in Z-axis rearing activity in both female and male *Winnie* mice compared to age- and sex-matched C57BL/6 controls (**Fig. 5E, 5E’**). Importantly, the rest period, defined as time spent without any detectable activity, was increased in *Winnie* mice of both sexes (females: *P* < 0.001; males: *P* < 0.01). Also, during the diurnal phase, the rest period was increased in female 5- and 15-week-old *Winnie* mice compared to age- and sex-matched C57BL/6 controls (*P* < 0.01) (**Fig. 5F, 5F’**). At 15 weeks of age, both absolute and body weight normalized forelimb grip strength in female and male *Winnie* mice showed no significant differences from age- and sex-matched C57BL/6 control mice (**Supplementary Fig. 4**).

**Figure 5.**
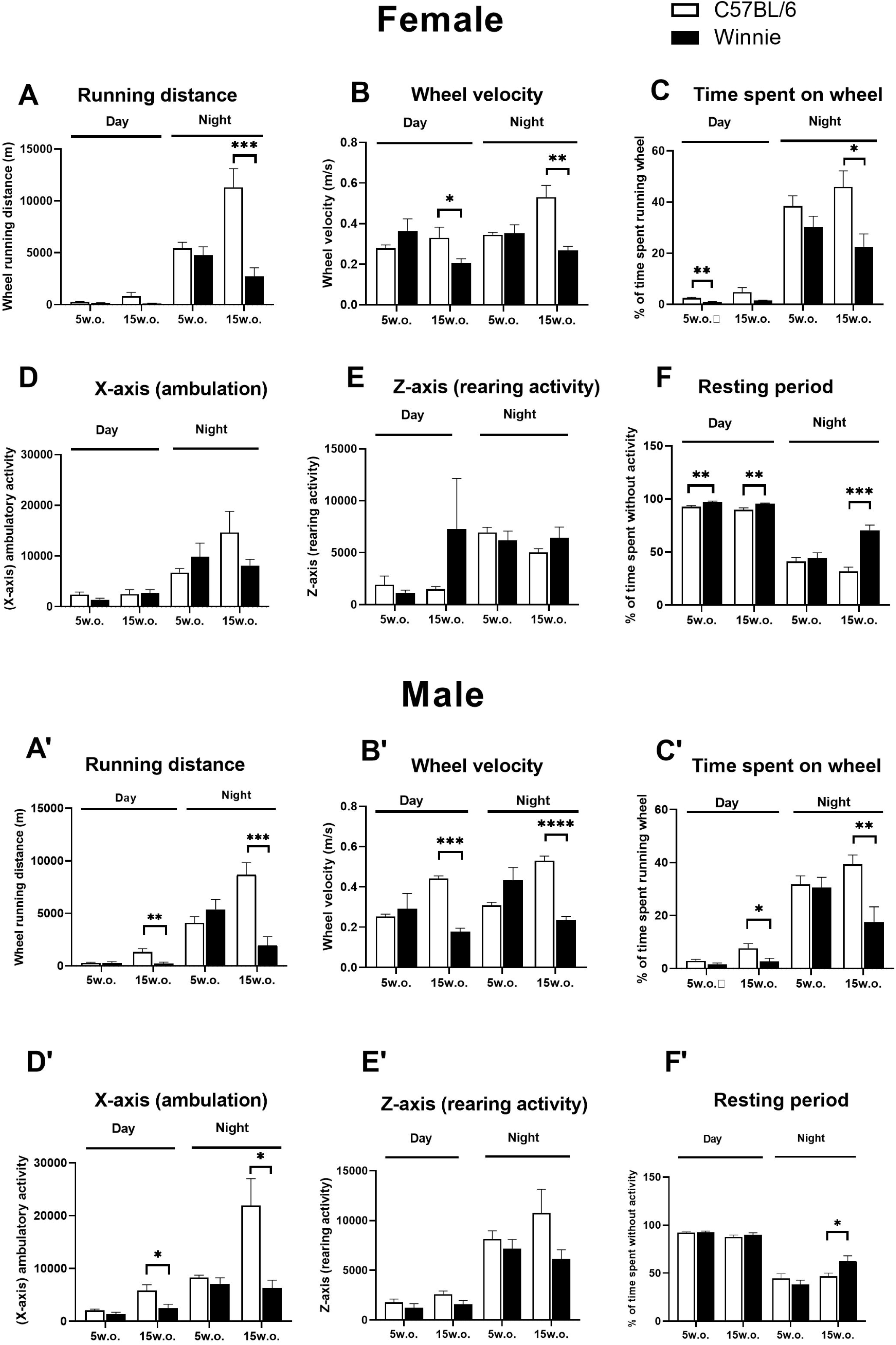
Promethion metabolic cage analysis for physical parameters in *Winnie* mice compared to C57BL/6 controls. (**A-F, A’-F’**) Distance covered, running velocity, percentage of time spent on the wheel, X-axis horizontal movement, Z-axis rearing activity and percentage of time spent inactive (rest period) during day and night cycle of female and male *Winnie* mice compared to C57BL/6 controls at 5 and 15 weeks of age. w.o. = weeks old. Data is represented as a mean ± SEM. *P < 0.05, **P < 0.01, ***P < 0.001, ****P < 0.0001; n=8 mice/group.

### Association between intestinal inflammation, DAI, and wheel running activity in Winnie mice with chronic colitis

LCN-2 levels showed a significant negative correlation with wheel running distance in *Winnies* at 15 weeks in both sexes, females (R^2^ = 0.511, *P* < 0.01) and males (R^2^ = 0.5082, *P* < 0.01). LCN-2 levels also negatively correlated with wheel velocity in *Winnies* at 15 weeks of age, with a stronger correlation observed in males (R^2^ = 0.777, *P* < 0.0001) compared to females (R^2^ = 0.5087, *P* < 0.01). The correlations were consistently stronger for wheel velocity compared to wheel running distance, particularly in male *Winnie* mice. DAI levels showed a negative correlation with wheel running distance in *Winnies* at 15 weeks in females (R^2^ = 0.4594, *P* < 0.01) compared to males (R^2^ = 0.3844, *P* < 0.05). DAI levels showed a negative correlation with wheel velocity in *Winnies* at 15 weeks in both sexes, females (R^2^ = 0.4602, *P* < 0.01) and males (R^2^ = 0.616, *P* < 0.001) (**Fig. 6**).

**Figure 6.**
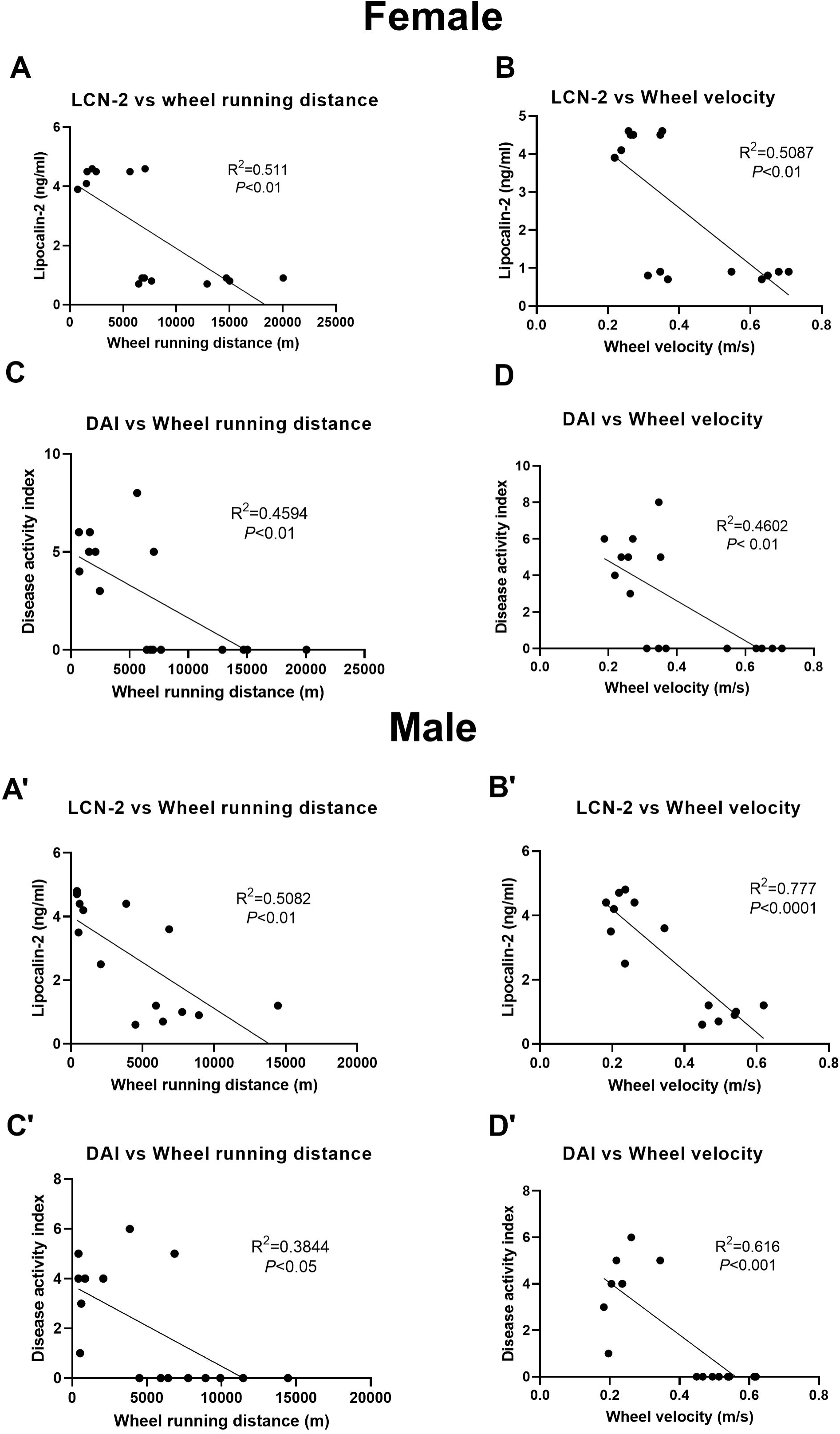
Linear regression analysis of Lipocalin-2 (LCN-2) levels and voluntary wheel running parameters in *Winnie* mice compared to C57BL/6 controls at 15 weeks. (**A, A’**) Correlations between LCN-2 levels and wheel running distance in females and males, respectively. (**B, B’**) Correlations between LCN-2 levels and wheel velocity in females and males, respectively. (**C, C’**) Correlations between DAI and wheel running distance in females and males, respectively. (**D, D’**) Correlations between DAI and wheel velocity in females and males, respectively. *P < 0.05, **P < 0.01, ***P < 0.001; n=14 mice/group. R^2^: coefficient of determination.

## Discussion

Muscle wasting is an extra-intestinal complication of IBD, characterized by a progressive and generalized loss of muscle mass and function. Recently, it has been reported that more than one-third of IBD patients suffer from significant muscle loss.^22^ This muscle wasting increases the risk of functional decline, falls, fractures, and hospitalizations in IBD patients.^23^ The substantial heterogeneity in evaluation methods for sarcopenia in IBD patients further complicates the identification of the underlying pathophysiology of IBD-associated muscle changes. Therefore, it is essential to develop animal models that feature chronic colitis-associated muscle wasting to better understand the mechanisms and progression of these muscle alterations in IBD.

Current animal models often fail to capture the chronic, progressive nature of IBD-related muscle changes, and the relationship between disease activity, inflammation, and muscle function, is not fully elucidated. This study aimed to characterize skeletal muscle phenotype and functional alterations in the *Winnie* mouse model, which spontaneously develops chronic colitis closely resembling human IBD,^1,8,24^ to address these gaps and provide insights into the muscle-gut axis in this condition. The present study provides a characterization of muscle phenotype and function in the *Winnie* mouse model of chronic colitis. We chose two specific time points to capture the transition from pre-disease to active disease states: 5–6 weeks (pre-inflammatory stage) and 15–16 weeks (active colitis stage). While more severe colitis develops in later stages (20-25 weeks), we focused on these earlier timepoints to investigate the initial impacts of colitis on muscle condition. This approach allowed us to examine how muscle changes correlate with the pre-onset and progression of colitis.

This study demonstrates that the *Winnie* mouse model of chronic colitis develops skeletal muscle alterations paralleling the progression of intestinal inflammation and a high disease activity at 15 weeks. These changes are characterized by reduced muscle mass and decreased fibre size, with functional implications primarily manifesting as reduced physical activity. A bidirectional relationship exists between muscle wasting and reduced activity. The muscle changes observed (reduced mass and fibre size) likely result from a combination of factors including chronic inflammation and reduced physical activity. These muscle changes, in turn, may further compromise the mice’s ability to engage in physical activity, creating a potential feedback loop (**Fig. 7**). This selective loss of muscle tissue highlights the catabolic nature of chronic inflammation and its specific impact on skeletal muscle. The absence of muscle changes at 5 weeks may suggest that a certain duration or threshold of inflammatory insult is necessary before detrimental effects on muscle health become evident. Our findings reveal a complex interplay between the progression of colitis and alterations in skeletal muscle, with implications for understanding muscle wasting and functional changes in IBD patients.

**Figure 7.**
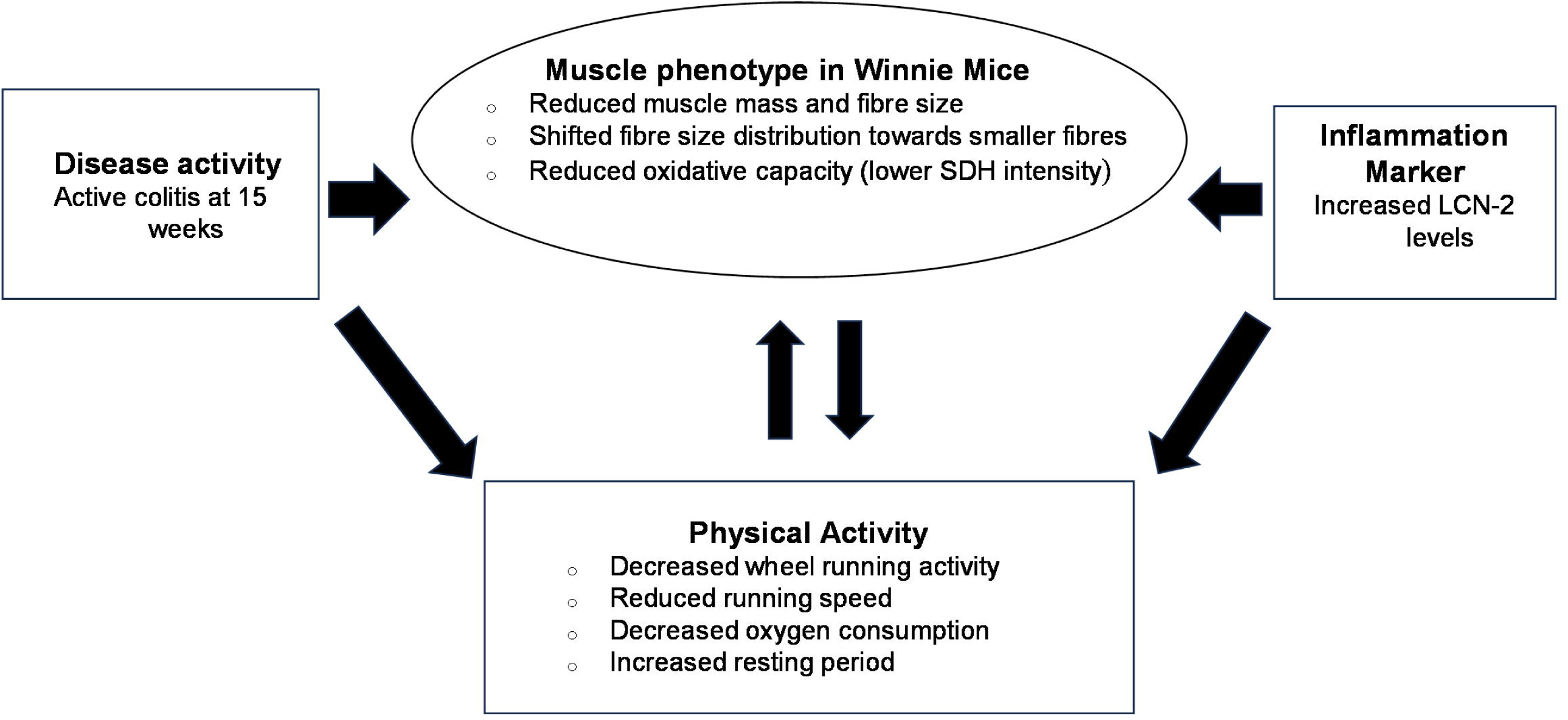
Schematic representation of muscle phenotype and associated changes in the *Winnie* mouse model of chronic colitis. This figure illustrates the interrelationships between disease activity, muscle changes, physical activity, and inflammation in *Winnie* mice with chronic colitis. The central box depicts the key muscle phenotype characteristics, including reduced muscle mass and fibre size and shifted fibre size distribution. Surrounding boxes show the associated changes in disease activity, physical activity, and inflammation markers.

Given that the degree of intestinal inflammation in genetically susceptible *Winnie* mice can vary between animal facilities due to environmental factors, particularly microbiota,^9,25^ we characterized the disease activity and extent of intestinal inflammation in our *Winnie* mouse colony under our specific experimental conditions. Our observations revealed that the *Winnie* mouse model exhibits a progressive development of colitis symptoms.

By 15 weeks of age, *Winnie* mice demonstrated significant differences compared to controls, including severity of colitis (characterized by reduced body weight, diarrhea, and rectal complications) and high intestinal inflammation as indicated by elevated lipocalin-2 (LCN-2) levels. LCN-2 is a protein that exhibits increased expression in the colonic epithelium and neutrophilic granulocytes.^26^ LCN-2 plays multiple roles in the intestinal inflammatory response, contributing to both local tissue reactions and systemic immune activation.^19^ Elevated intestinal inflammation observed in *Winnie* mice with chronic colitis was consistent with findings from mouse models of IBD that have established the role of LCN-2 in intestinal inflammation and gut bacterial dysbiosis.^19,27,28^ These elevated LCN-2 levels show a strong correlation with well-established markers of inflammation and disease activity in IBD, suggesting that LCN-2 plays a significant role in the underlying pathophysiological mechanisms of IBD.^26,29^

The absence of these symptoms in age-matched C57BL/6 control mice confirms that these changes are specific to the *Winnie* genotype and not attributable to normal aging processes. Importantly, this disease progression coincided with notable changes in muscle phenotype and function. Preliminary *in vivo* observations revealed reduced lean mass without corresponding changes in fat mass in *Winnie* mice. This altered body composition mirrors reports from studies of IBD patients, suggesting that our model recapitulates key aspects of the human disease phenotype.^30^

We selected locomotory muscles, TA and SOL, as representatives of fast-twitch and slow-twitch muscles, respectively. The mouse SOL has been reported to express the closest molecular resemblance to several human skeletal muscles.^31^ In 15-week-old *Winnie* mice, we observed decreased mass in both SOL and TA muscles. This finding corroborates the study by Bryant et al., which reported up to 60% reduction in muscle mass in IBD patients.^32^

The absence of significant differences at 5 weeks suggests that muscle fibre atrophy is not an early feature of colitis in the *Winnie* model, aligning with the progression of intestinal inflammation in these mice. The significant reduction in muscle fibre size by 15 weeks in both SOL and TA muscles indicates that chronic colitis leads to widespread muscle atrophy affecting both slow-twitch and fast-twitch fibres. The more pronounced shift in fibre size distribution in TA muscles suggests that fast-twitch fibres may be more susceptible to atrophy in this model of chronic colitis. Several studies have indeed shown that fast-twitch fibres, which are predominant in the TA muscle, can be more susceptible to certain types of stress or damage compared to slow-twitch fibres, which are predominant in the SOL muscle.^33,34^ It has been reported that the heightened vulnerability of fast-twitch muscles to inflammation-induced wasting could be attributed to their greater reliance on glycolytic metabolism and higher protein turnover rates, which could make them more susceptible to the catabolic effects of inflammation.^33^

The observed muscle fibre atrophy corroborates the previously noted decreases in muscle mass, providing a cellular basis for the macroscopic changes in muscle size. Importantly, the general muscle architecture of SOL and TA muscle fibres remained intact, and the absence of fibre attrition suggests that *Winnie* mice do not exhibit a phenotype of severe muscle damage. However, the shift in muscle fibre size distribution towards smaller sizes indicates preferential atrophy or loss of larger muscle fibres. This preferential atrophy of larger fibres suggests a potential metabolic component to the muscle wasting process, which warrants further investigation The robust correlation between intestinal inflammation (as measured by LCN-2 levels) and muscle wasting parameters highlights the intricate relationship between gut health and systemic muscle function in IBD. It has been shown that LCN-2 can have systemic implications and be detected in circulation during active inflammatory conditions.^28^ The observed correlations between LCN-2 levels and muscle parameters indicate a possible pathway (release of inflammatory mediators, alterations in gut permeability, or changes in the gut microbiome) by which intestinal inflammation could affect distant tissues like skeletal muscle. Recent studies are exploring how locally produced LCN-2 might influence distant tissues in IBD.^35^ The strong association between LCN-2 levels and muscle wasting parameters suggests that LCN-2 could potentially serve as a biomarker for predicting muscle changes in IBD. This aligns with previous literature showing that LCN-2 influences muscle phenotype in mouse model of muscular dystrophy.^36^ Interestingly, we found that these correlations were stronger for the TA muscle compared to the SOL, suggesting potential muscle-specific responses to inflammation. This emphasizes the need for a holistic approach to IBD management that considers both intestinal and extra-intestinal manifestations of the disease.

The observed correlations between Disease Activity Index (DAI) and various muscle measurements highlight the systemic impact of chronic intestinal inflammation on skeletal muscle health. The results suggest sex-specific responses to chronic colitis in terms of muscle wasting. In females, the SOL muscle appears more affected, while in males, both muscles show similar correlations with disease activity. The SOL and TA muscles show different degrees of correlation with disease activity, suggesting that muscle fibre type may influence susceptibility to inflammation-induced atrophy. Despite variations in correlation strength, all muscle parameters negatively correlated with disease activity in both sexes, indicating a consistent detrimental effect of colitis on muscle health. The correlations suggest that systemic inflammation associated with colitis may directly impact muscle mass and fibre size, possibly through altered protein metabolism or increased catabolism. These findings in the *Winnie* mouse model align with clinical observations of muscle wasting in IBD patients, supporting the model’s translational value. The strong correlations between disease activity and muscle parameters suggest that treatments targeting inflammation might also benefit muscle health in IBD. Further investigation into the molecular pathways linking intestinal inflammation to muscle-specific atrophy could reveal new therapeutic targets for preserving muscle mass and function in IBD. While these correlations are strong, they do not prove causation. Interventional studies would be needed to establish a direct causal link between disease activity and muscle wasting. However, these results underscore the systemic nature of IBD, highlighting the importance of considering extra-intestinal manifestations like muscle wasting in the comprehensive management of the disease.

The reduction in both muscle mass and fibre size with increasing disease activity may contribute to the observed decreases in physical activity and function in colitis models. We assessed locomotory ability in *Winnie* mice using a voluntary wheel running paradigm implemented in the Promethion metabolic cage system. Voluntary wheel running offers several advantages as an assessment tool. Notably, it elicits running patterns similar to the natural locomotor behaviour of mice under non-stressed conditions, aligning with the animal’s intrinsic rhythmicity.^37^ This approach allows for a more naturalistic evaluation of physical capacity compared to forced exercise protocols. Our analysis of physical performance parameters revealed decrease in both the quantity (wheel running distance and time) and quality (velocity) of physical activity in *Winnie* mice. The maintenance of horizontal movement in females, despite reduced wheel running, suggests sex-specific muscle adaptations or compensatory mechanisms that preserve muscle function for essential movements, possibly through hormonal influences. Increased inactivity time implies that the colitis mice spent more time being sedentary or inactive, which is a manifestation of reduced physical capacity and functional limitations. Consistently, impaired physical activity has been reported in IBD patients.^38,39^

The significant negative correlations between LCN-2 levels and wheel running parameters (distance and velocity) suggest a strong link between intestinal inflammation and reduced physical activity in this model of chronic colitis. The stronger correlation between LCN-2 and wheel velocity in males indicates that intestinal inflammation may have a more pronounced effect on the intensity of physical activity in male mice. The negative correlations between DAI and wheel running parameters demonstrate that overall disease severity impacts physical activity, supporting the systemic nature of the disease’s effects. The stronger correlations observed with wheel velocity compared to distance for both LCN-2 and DAI indicate that these markers of inflammation and disease activity may be particularly associated with the ability to perform high-intensity physical activity. These findings support the hypothesis that both localized intestinal inflammation (as indicated by LCN-2) and overall disease activity (as measured by DAI) are associated with reduced physical performance in *Winnie* mice with chronic colitis. Long-term changes in *Winnie* mice with chronic colitis reflect a combination of disease activity and muscle adaptations. The data provide quantitative evidence for the relationship between inflammatory markers, disease activity, and functional outcomes, strengthening the connection between gut health and overall physical well-being in this model. The findings suggest that monitoring both LCN-2 levels and DAI might provide insights into physical function impairments in chronic inflammatory conditions, which could have translational implications for assessing and managing IBD patients.

This explanation emphasizes that in *Winnie* mice, as in human IBD, the relationship between disease activity, muscle health, and physical activity is multifaceted and bidirectional, rather than a simple cause-and-effect scenario. This complex interplay underscores the importance of considering both direct disease effects and secondary muscle changes when studying physical impairment in chronic colitis.

The negative correlation between LCN-2 levels and physical activity in mice is consistent with some human studies. For instance, DeFilippis et al. (2016) found that elevated fecal calprotectin (another marker of intestinal inflammation) was associated with reduced physical activity in IBD patients.^40^ The correlation between DAI and reduced physical activity in *Winnie* mice at 15 weeks aligns with clinical studies showing that higher disease activity in IBD is associated with reduced physical function. Ploeger et al. (2012) reported that IBD patients with active disease had lower exercise capacity compared to those in remission.^41^ The strongest and most significant relationships are observed in females between DAI, LCN-2, and both wheel running distance and velocity. The stronger correlations with wheel velocity compared to distance in mice suggest that inflammation and disease activity might particularly affect high-intensity activities. In human studies, Tew et al. (2016) found that IBD patients had reduced anaerobic threshold during exercise, which could parallel the reduced high-intensity activity in mice.^42^ While LCN-2 is not commonly used in human IBD studies, other markers like fecal calprotectin or C-reactive protein have been associated with physical function in IBD patients.^43^ Future studies could investigate the mechanisms by which intestinal inflammation and disease activity impact physical activity, potentially involving factors such as fatigue, muscle metabolism, or neuroinflammatory pathways.

Interestingly, while wheel-running activity was reduced in *Winnie* mice with active colitis, forelimb grip strength remained unaltered. The maintenance of grip strength in *Winnie* mice at 15 weeks suggests that the colitis model does not impair maximal force production in the muscles involved in gripping. This discrepancy between different functional measures might indicate that the neuromuscular junctions and motor unit recruitment patterns remain efficient for short, intense actions but may be compromised for sustained or repetitive movements. The relatively intermittent and short-duration nature of grip strength tasks, compared to sustained activities like wheel running, may allow for more effective compensatory mechanisms to maintain performance. The contrast between preserved grip strength and reduced wheel running activity points to a more pronounced effect on muscle endurance rather than maximal strength.

Our earlier observations in *Winnie* mice showed normal food intake, effectively ruling out malnutrition and decreased appetite as contributing factors to muscle loss. These findings parallel observations in UC patients, where the incidence of nutrient deficiencies is less compared to other forms of IBD, however, weight loss is common among them.^44,45^ The absence of nutrient deficiencies in our model, despite evident muscle mass loss, highlights the significant role of chronic inflammation in muscle changes in IBD. However, it’s important to note that muscle loss in chronic inflammatory conditions like IBD is complex and likely involves multiple mechanisms, including systemic inflammation, altered metabolism, and reduced physical activity. Our findings contribute to understanding this multifaceted process, particularly emphasizing the potential primary role of chronic inflammation. This accentuates the potential importance of anti-inflammatory therapies for managing gut symptoms and preserving muscle health.

Fatani et al. (2023) proposed a classification system for muscle damage in IBD patients: myopenia (low muscle mass), pre-sarcopenia (low muscle strength), and sarcopenia (low muscle mass and strength).^3^ Based on this classification and our findings, we conclude that the *Winnie* mouse model exhibits a myopenic phenotype, characterized by low muscle mass/size but preserved muscle strength. This nuanced characterization provides valuable insights into the progression of muscle changes in IBD, suggesting that interventions to maintain muscle mass may be beneficial even before functional deficits become apparent.

The progressive nature of muscle changes in the *Winnie* model, correlating with the onset and progression of colitis, provides a valuable framework for understanding the development of muscle deterioration in IBD patients. The preservation of certain muscle functions (grip strength) alongside reduced physical activity (wheel-running) suggests that early interventions targeting muscle preservation might be beneficial, even before overt functional deficits are observed.

While our study provides comprehensive insights into muscle changes in chronic colitis, it has limitations. The use of a single mouse model may not capture the full spectrum of IBD-associated muscle changes. Future studies may explore these phenomena in other IBD models. The preservation of grip strength despite muscle changes suggests complex adaptations that warrant further investigation. Additionally, longitudinal studies in IBD patients, correlating muscle changes with disease progression, would further validate the translational relevance of our findings.

## Conclusions

This study demonstrates that the *Winnie* mouse model of chronic colitis develops skeletal muscle alterations paralleling the progression of intestinal inflammation. Study provides novel insights into the temporal progression of muscle wasting in the *Winnie* mouse model of IBD. Our findings validate the *Winnie* mouse as a valuable pre-clinical model for investigating IBD-associated muscle atrophy. These findings underscore the systemic nature of IBD and highlight the potential importance of monitoring and maintaining muscle function in this condition. Also, it can be inferred that the muscle-wasting effects of chronic intestinal inflammation are not uniform across all muscle functions, but represent a nuanced process affecting functional capacity in specific ways. Future studies should focus on translating these findings to IBD patients, exploring the time course of muscle changes in disease progression, and investigating strategies to preserve muscle mass and function in the context of chronic intestinal inflammation.

## Acknowledgments

Shilpa Sharma thanks the Australian Government Research Scholarship and a Seed Grant from the Australian Institute for Musculoskeletal Science (AIMSS).

---

**Supplementary Figure 1.** Image showing the anal region of C57BL/6 and Winnie mice at 15 weeks old. The prominent rectal prolapse in the *Winnie* mouse, indicated by the protruding rectal tissue (arrow).

**Supplementary Figure 2. Lean and fat mass in *Winnie* mice at 15 weeks compared to age-matched C57BL/6 controls.** (**A**) Lean mass and (**B**) fat mass in *Winnie* mice at 15 weeks compared to age-matched C57BL/6 controls. w.o. = weeks old. Data is represented as a mean ± SEM. **P < 0.01; n=12 mice/group.

**Supplementary Figure 3. Linear regression analysis of Disease Activity Index (DAI) and muscle mass and size in *Winnie* and C57BL/6 mice at 15 weeks.** (**A, A’**) Correlations between DAI and SOL mass in females and males, respectively. (**B, B’**) Correlations between DAI and TA mass in females and males, respectively. (**C, C’**) Correlations between DAI and SOL fibre size in females and males, respectively. (**D, D’**) Correlations between DAI and TA fibre size in females and males, respectively.

**Supplementary Figure 4. The forelimb grip strength in *Winnie* and C57BL/6 mice at 15 weeks.** Absolute and relative forelimb grip strength of (**A, B**) female and (**A’, B’**) male *Winnie* mice at 15 weeks compared to C57BL/6 controls.

