## Supplementary figures and images for "Advancements in Inflammation Parallels Myopenia in *Winnie* Mice Model of Spontaneous Chronic Colitis"

### Supplementary Figure 1

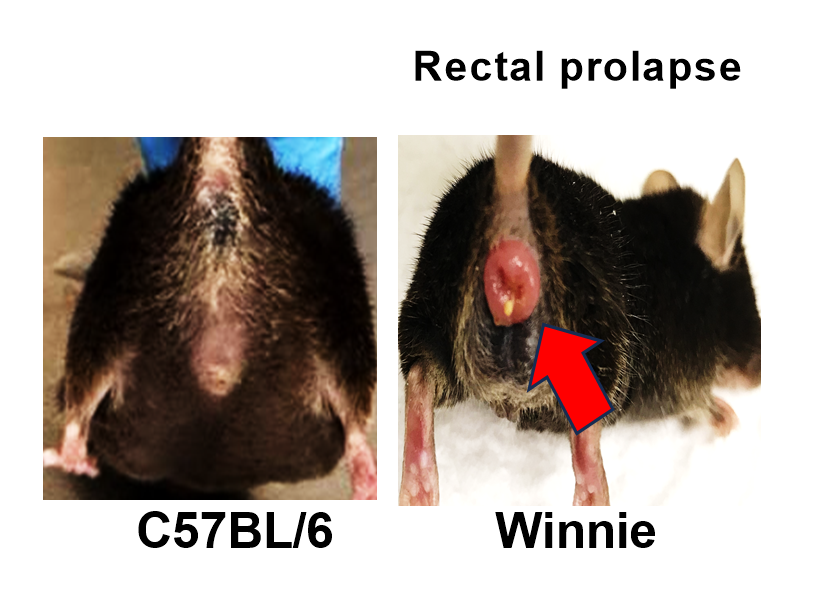

### Supplementary Figure 2

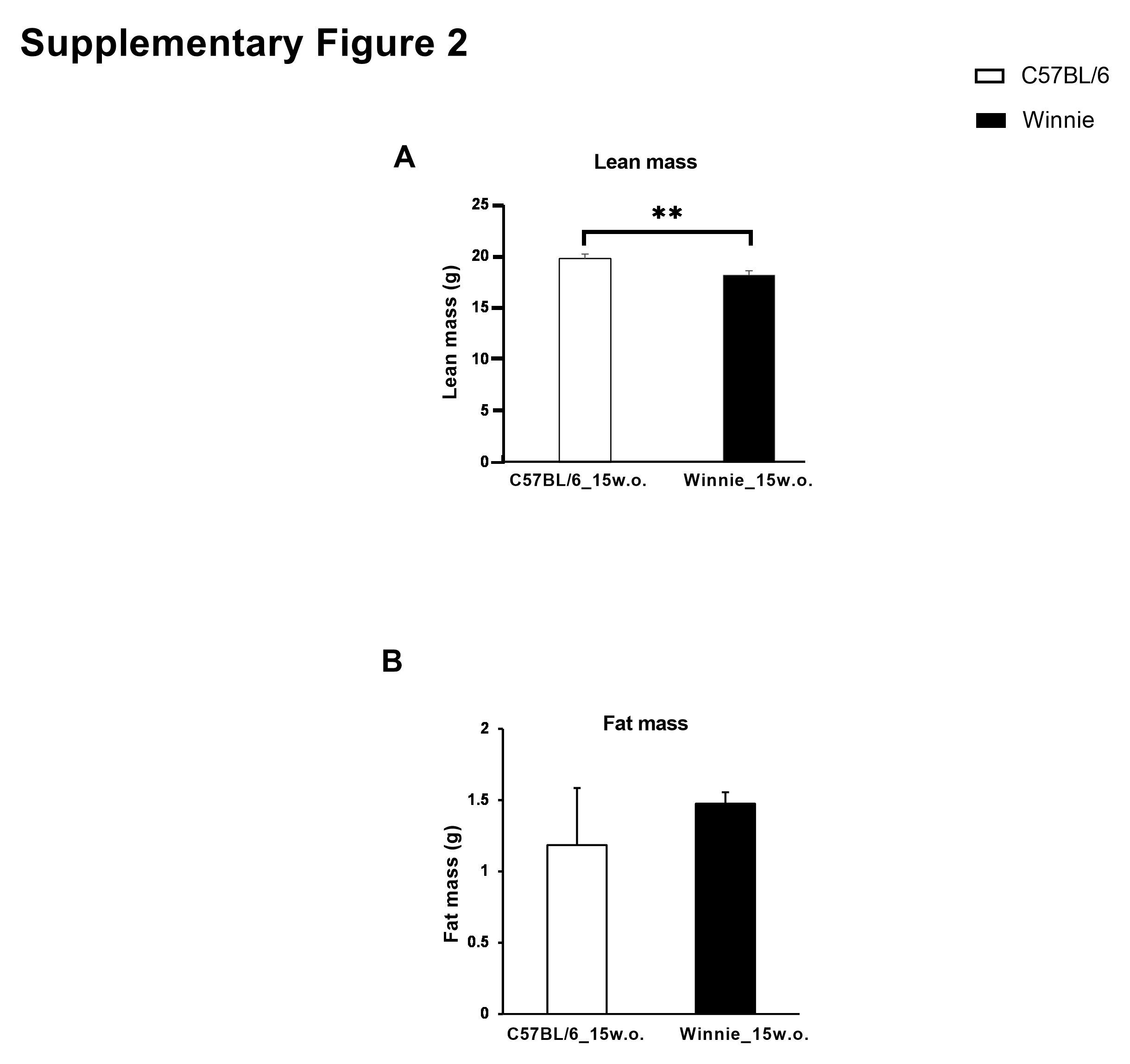

### Supplementary Figure 3

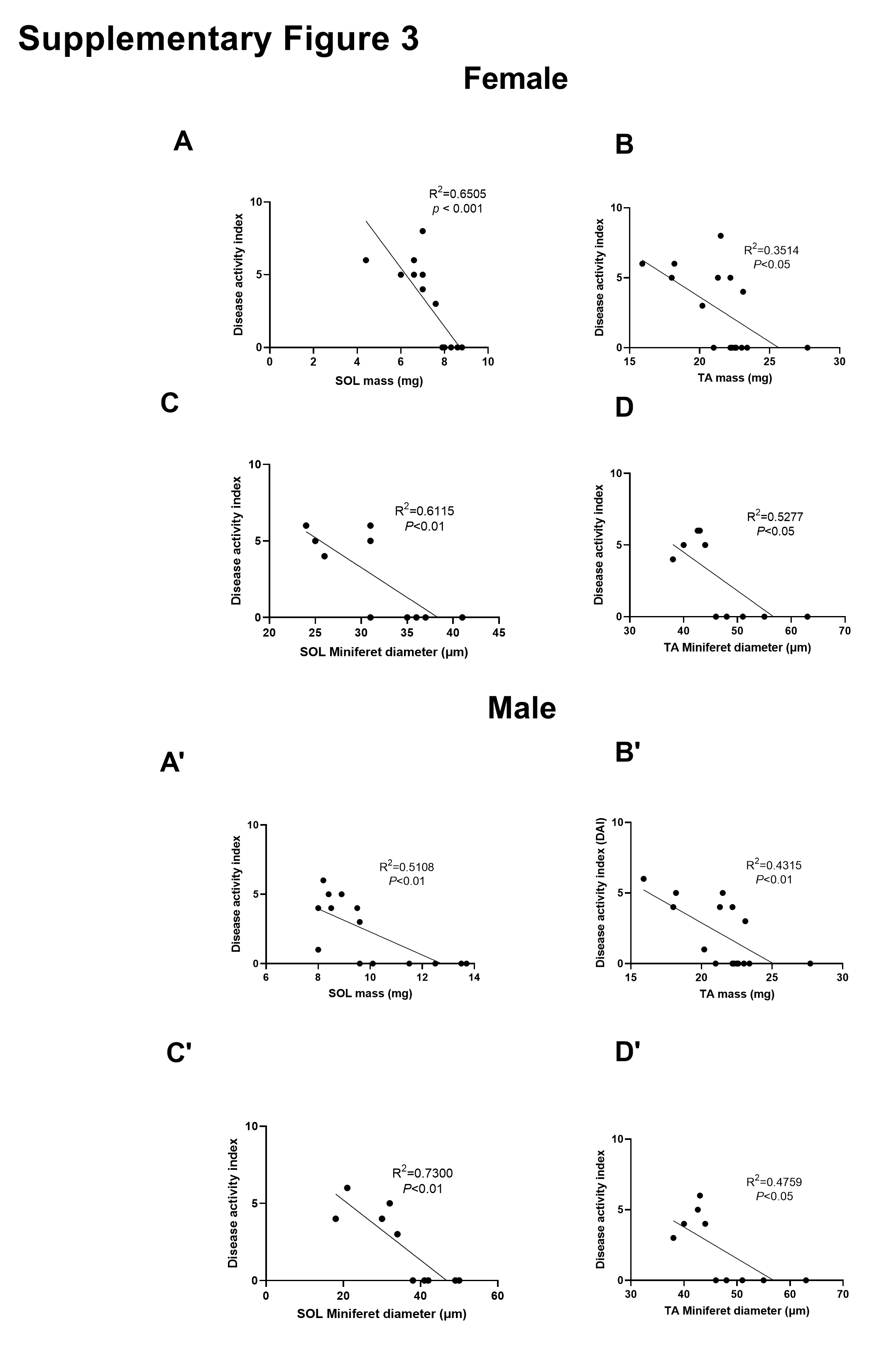

### Supplementary Figure 4

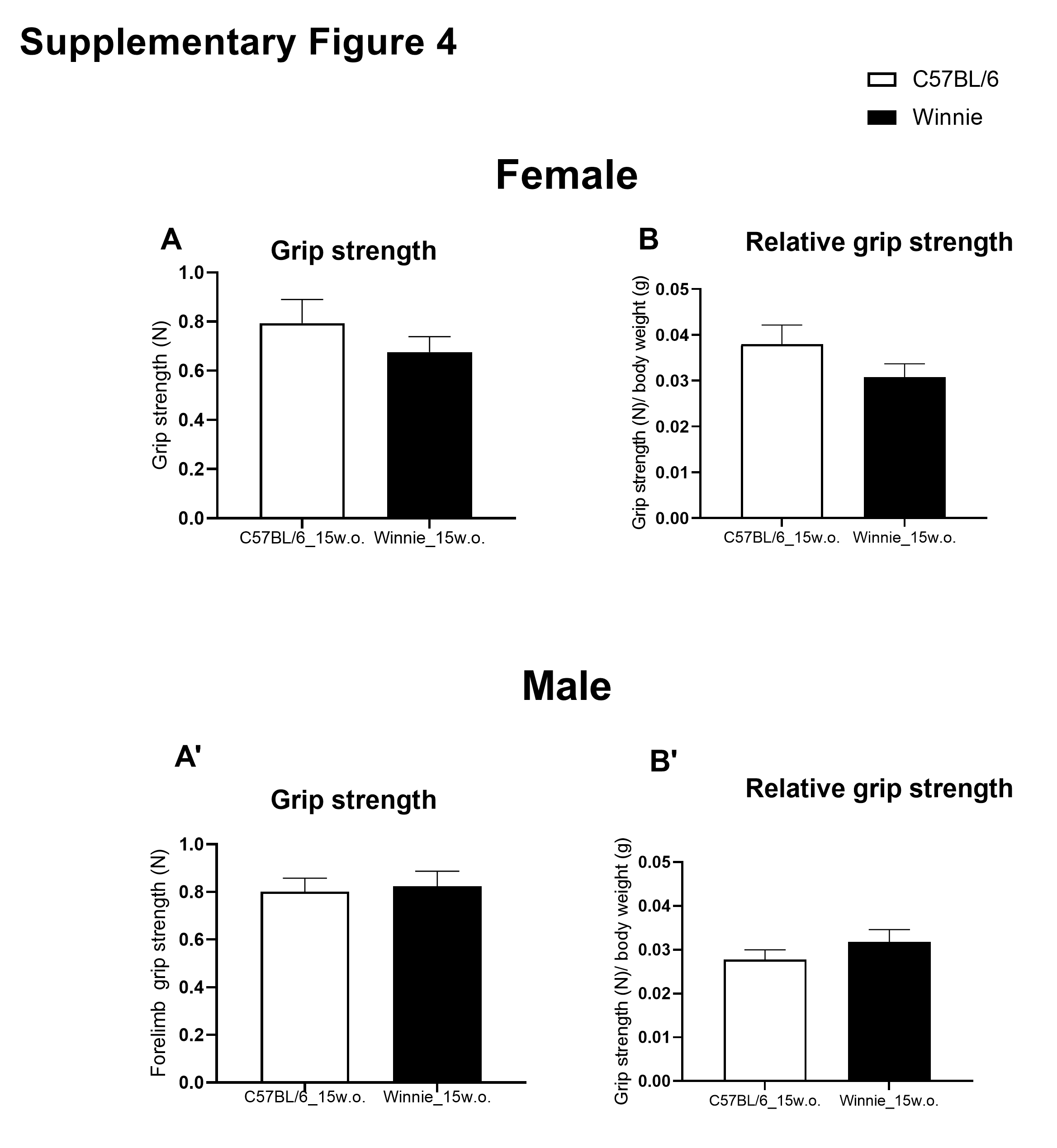
